# IMMUNE GENE REGULATION IN THE GUT DURING METAMORPHOSIS IN TWO HOLO- VERSUS A HEMIMETABOLOUS INSECT

**DOI:** 10.1101/2022.06.26.497637

**Authors:** Christin Manthey, Paul R. Johnston, Jens Rolff

## Abstract

During complete metamorphosis, holometabolous insects remodel their entire anatomy, including the gut and must control their microbiota to avoid infectious disease. High activity of antimicrobial peptides and proteins in the gut during metamorphosis has been best described in several Lepidoptera. The immune system of the dipteran *Drosophila melanogaster* also controls the number of bacteria during metamorphosis. However, little is known about the regulation of immune genes during the larval–adult moult in Hemimetabola which undergo less drastic metamorphic changes. Different patterns of immune effector expression during metamorphosis were shown in a study comparing the lepidopteran *Galleria mellonela* and the orthopteran *Gryllus bimaculatus,* where *G. mellonella* showed a strong up-regulation of antimicrobial peptides in the gut at the larval-pupal moult but not *G. bimaculatus.* Whether these findings reflect general patterns within holometabolous versus hemimetabolous insects remains unclear. Using RNAseq, we compare the expression of immune effector genes in the gut during metamorphosis in two holometabolous (*Calliphora vicina* and *Tenebrio molitor*) and a hemimetabolous insect (*Pyrrhocoris apterus).* We found high read count abundances of differentially expressed immune effectors in the gut at the larval-pupal moult in *C. vicina* and *T. molitor;* no such high abundances were observed at the larval-adult moult in *P. apterus.* Our findings confirm that only complete metamorphosis elicits a prophylactic immune response as an adaptive response in holometabolous insects, which controls the microbiota during gut replacement.

## Introduction

Complete metamorphosis is considered a key trait that explains the incredible diversity of insects (Nicholson, Ross, & Mayhew, 2014). The drastic reconstructions in the pupal stage in insects with complete metamorphosis (Holometabola) (Hall & Martín-Vega, 2019) allow for decoupling of growth and differentiation (Arendt, 1997, Rolff et al. 2019). It also gives the insect the unique opportunity to change the microbial composition throughout insect development (Manthey, Johnston, & Rolff, 2021), facilitating niche shifts between the larval and adult life stages (Hammer & Moran, 2019; Rolff, Johnston, & Reynolds, 2019).

However, the drastic reconstruction during complete metamorphosis puts the insect at a higher risk of infections. The physical barriers in the gut that avoid bacterial infections, like the peritrophic membrane in the midgut and the sclerotised cuticle of the fore- and hindgut, are broken down during complete metamorphosis. Hence, the insect must control its gut microbiota to avoid infectious disease, creating a dilemma; either the insect host eradicates and reestablishes its gut microbes from the environment, or the insect maintains beneficial symbionts while fighting pathogens. It is known that insects have a mixed-mode transmission of microbes (Ebert, 2013). Some beneficial symbionts are transmitted vertically and stored in specialised tissues during complete metamorphosis, while others are horizontally transmitted and taken up from the environment later. Insects evolved various strategies to ensure the transmission of beneficial symbionts throughout development. Stoll, Feldhaar, Fraunholz, & Gross (2012) showed vertical transmission of microbes via bacteriocytes in ant species. The relative number of bacteria-filled bacteriocytes increased strongly during complete metamorphosis. Maire et al. (2020) also showed a transmission of microbes via bacteriocytes in weevils by maintaining and relocating bacteriocytes during gut renewal in the pupa. Other specialised structures to transmit symbionts in insects are antennal glands (Kaltenpoth, Yildirim, Gürbüz, Herzner, & Strohm, 2012) and crypts (Kikuchi, Hosokawa, & Fukatsu, 2011).

Also, insects initiate immune responses to control their gut microbiota and avoid infectious diseases. When in contact with a pathogen, insects defend themselves using cellular and humoral immunity (Du Pasquier, 2001; Hultmark, 1993). Immediate reactions include the induction of proteolytic cascades, such as activating phenoloxidase that affects melanin formation (Zhao, Li, Wang, & Jiang, 2007). Among the induced effector molecules are antimicrobial proteins and peptides (AMPs).

It is likely that holometabolous insects evolved strategies to pre-emptively activate immune processes in the pupal gut to prevent infections during complete metamorphosis. Russell & Dunn (1991) described high activity of the antimicrobial protein lysozyme in the midgut lumen of the moth *Manduca sexta* (Lepidoptera) during complete metamorphosis. Lysozyme accumulated in the larval midgut epithelium and was subsequently released into to the gut lumen at the larval-pupal moult. Also, Russell & Dunn (1996) found that the pupal midgut of *M. sexta* contains a cocktail of antimicrobial proteins, including at least lysozyme, bactericidal activity against *Escherichia coli,* hemolin, and phenoloxidase. Induction of antimicrobial peptides in the gut prior to complete metamorphosis has also been described in other Lepidoptera, the silkworm *Bombyx mori* and the tobacco cutworm *Spodoptera litura* (Mai et al., 2017). Mai et al. (2017) found up-regulated lebocin, an AMP specific to Lepidoptera, in the midgut with a peak expression during the wandering stage. They also found that the ecdysteroid hormone 20-hydroxyecdysone (20E), which is known to control metamorphosis, regulates lebocin expression in the midgut. Nunes, Koyama, & Sucena (2021) confirm the link between the endocrine and immune systems to control the number of bacteria in the pupa of the fruit fly *Drosophila melanogaster* (Diptera). They found three AMPs (*drosomycin, drosomycin-like 2* and *drosomycin-like 5*) were differentially expressed at pupation irrespective of the presence of bacteria and regulated by 20E. This shows that co-option of immune effector gene expression by the 20E moulting pathway is not restricted to the Lepidoptera and may be a general phenomenon in the Holometabola.

In contrast, in insects with incomplete metamorphosis (Hemimetabola), the metamorphic changes are less drastic, and the gut microbiota stays relatively stable compared to holometabolous insects (Manthey et al., 2021). However, little is known about the regulation of immune genes during the larval-adult moult. Johnston, Paris, & Rolff (2019) were the first to show that hemimetabolous and holometabolous insects have different patterns of immune effector expression. They found a strong up-regulation of antimicrobial proteins and peptides and the transcription factor GmEts at the onset of pupation in the greater wax moth *Galleria mellonella* (Lepidoptera), but no such up-regulation at the larval–adult moult in the hemimetabolous cricket *Gryllus bimaculatus* (Orthoptera). *G. mellonella* showed peak expression of lysozymes and three AMPs coinciding with delamination of the larval gut. However, whether these findings reflect general patterns within holometabolous and hemimetabolous insects remains unclear.

Here we use RNA-seq to compare the temporal dynamics of immune effectors expression in the gut at the larval-pupal moult in three other orders of insects, two holometabolous insects, the blow fly *Calliphora vicina* (Diptera) and the mealworm beetle *Tenebrio molitor* (Coleoptera) and at the larval-adult moult in a hemimetabolous insect, the firebug *Pyrrhocoris apterus* (Hemiptera). According to Johnston et al. (2019), we hypothesized that an immune effector expression would be iniatited at the onset of complete metamorphosis in the two holometabolous insects. By contrast, given that the gut does not undergo drastic reconstruction during incomplete metamorphosis and in the absence of infection, we expect no immune effector induction in the hemimetabolous insect during the final moult..

## Material and Methods

### *Tenebrio molitor* rearing and sampling

*Tenebrio molitor* (mealworm beetle) larvae were purchased from a commercial supplier (Der Terraristikladen, Düsseldorf, Germany) and used to establish a laboratory colony at the Freie Universität Berlin. The mealworm beetles were held in fauna-boxes (Reptilienkosmos, Viersen, Germany) with a 14 L: 10 D cycle and 60 ± 5% humidity at 25 ± 1°C. They were fed wheat bran supplemented with carrot and water *ad libitum*. The mealworm beetles were sampled in the final durint six sampling stages:

I. Final moult stage, a freshly moulted, large last instar larva with a still translucent white cuticle maximum of one hour after the final moult.
II. Little movement stage, a larva 1-2 d before pupariation resting on the substrate, but responding to pinch grips.
III. No movement stage, a larva 12 h before puparation resting on the substrate and not responding to pinch grips.
IV. Pupa after 1 – 6 h, a white soft pupa 1-6 h after pupation.
V. Pupa after 12-20 h, a light brown beige pupa 12-20 h after pupation.
VI. Pupa after 24-48 h, a light brown beige pupa with clearly black eyes 24-48 h after pupation.

RNA was isolated from dissected guts as described below and used to create three independent replicate pools per stage, each representing five individual insects, resulting in 18 sample pools.

### *Calliphora vicina* rearing and sampling

A laboratory colony of *Calliphora vicina* (urban bluebottle blowfly) was established with insects purchased from a commercial supplier (Reptilienkosmos, Viersen, Germany) and reared at the Freie Universität Berlin. The blowflies were held in insect gauze cages (BugDorm, Taichung, Taiwan) in an incubator with a 14 L: 10 D cycle and 60 ± 5% humidity at 25 ± 1°C. They were fed a diet of milk powder, sugar (ratio 3:1) and water ad libitum. Additionally, calf’s liver was offered for oviposition. Blowflies were checked daily to determine their development. The larvae developed over three larval stages, with the third larval instar being divided into the feeding and post-feeding stages. The first and second larval stages lasted one day, and the final instar larva lasted five to seven days. The blowflies were sampled at the onset of complete metamorphosis in the post-feeding and pupal stages. The sampling stages in the post-feeding and pupa were specified as follows:

I. Post-feeding larval stage, a still actively crawling but non-feeding third instar larva.
II. Pre-pupal stage, a white, contracted larva responding to pinch grips; just before the transition to white puparium; shiny white and soft cuticle.
III. Pupal stage at the onset, a white and motionless puparium ready for pupariation; cuticle was dull white and dried (unlike stage II).
IV. Pupa after 1 h, a medium brown tubule emerged one hour after stage III; the cuticle was slightly more hardened than in stage III.
V. Pupa after 4-6 h, a reddish-brown cryptocephalous pupa, 4-6 h after stage III; the puparium was fully hardened; beginning of larval-pupal apolysis.
VI. Pupa after 8-12 h, a blackish-brown pupa 8-12 h after stage III; larval pupal apolysis.

RNA was isolated from dissected guts and used to create three independent replicate pools per stage, each representing five individual insects, resulting in 18 sample pools.

### *Pyrrhocoris apterus* rearing and sampling

A laboratory culture of *Pyrrhocoris apterus* (firebug) was established with insects collected from *Tilia cordata* (small-leaved linden) trees at three locations in Berlin (see table 1, supplement) in April 2021. Only firebugs not parasitized by mites were collected. The firebugs were reared in fauna-boxes (Reptilienkosmos, Viersen, Germany) at the Freie Universität Berlin with a 14 L: 10 D cycle at room temperature (21 ± 1°C). They were fed *T. cordata* seeds and water ad libitum via cotton plugged tube. After oviposition, the eggs were held in plastic boxes covered with a gauze until hatching. Firebugs were reared individually from the fourth instar in small plastic boxes covered with gauze and checked daily to determine their development. The larvae developed over five instars within a maximum of 40 days. The fifth instar lasted ten days. Five stages in the last instar larva and the adult were sampled from the F1-generation and specified as follows:

I. Final instar larva, a freshly moulted, fifth instar larva that is still decoloured and entirely reddish-orange.
II. Final instar after 4-5 days, a fifth instar larva four to five days after stage I.
III. Final instar after 9 days, a last instar larva nine days after stage I.
IV. Final instar after 9.5 days, a last instar larva nine and a half days after stage I.
V. Adult, a freshly eclosed adult maximum four hours after the imaginal moult.

Isolated RNA from guts was used to create three independent replicate pools per stage, each representing five individual insects, resulting in 18 sample pools.

### RNA isolation and library preparation

Insect guts were dissected with dissecting utensils sterilised with ethanol. The dissected guts were rinsed in distilled water, and RNA was extracted from the guts with Trizol. They were homogenized in 1ml Trizol (Sigma, Taufkirchen, Germany) with two sterile 3 mm beads (Qiagen, Hilden, Germany) and a bead device at 20 Hz for 3 min. According to the manufacturers’ instructions, total RNA was recovered using chloroform phase separation and isopropyl alcohol precipitation. RNA concentrations were measured using the Qubit™ RNA High Sensitivity Kit and a Qubit 4 fluorometer (ThermoFisher, Schwerte, Germany). Equal quantities of the samples were used to create independent replicate pools for each stage. RNA pools were purified by incubating the samples with TurboDNase (© Ambion) for 30 min at 37°C and cleaned up with the RNeasy MiniElute cleanup kit (Qiagen, Hilden, Germany) following the manufacturers’ instructions. The concentrations of the pooled RNA samples were measured using the Qubit™ RNA HS Assay Kit and a Qubit™ 4 fluorometer (ThermoFisher, Schwerte, Germany). Samples from *Tenebrio molitor* and *Calliphora vicina* were qualified with the High Sensitivity RNA ScreenTape Assay Kit and the Agilent 4200 TapeStation system (Agilent, Santa Clara, United States) and *Pyrrhocoris apterus* samples with the Agilent RNA 6000 Pico Kit on a BioAnalyzer 2100 (Agilent, Santa Clara, United States). *T. molitor, C. vicina* and *P. apterus* libraries were prepared using the NEBNext^®^ Ultra II Directional RNA Library Prep Kit (New England Biolabs, Ipswich, United States) at the Berlin Center for Genomics in Biodiversity Research (BeGenDiv). Library qualities were assessed with the High Sensitivity D1000 ScreenTape Assay Kit and the Agilent 4200 TapeStation system (Agilent, Santa Clara, United States). The libraries of all three insect species were sequenced on NovaSeq 6000 at the Institute of Clinical Molecular Biology (IKMB) in Kiel for 300 cycles to yield 15–41 million 150-bp read pairs per library (mean 28 million).

### *De novo* assembly and annotation

Assemblies for both species were produced using Trinity v. 2.8.4 (Haas et al., 2013), incorporating quality and adapter filtering via Trimmomatic (Bolger, Lohse, & Usadel, 2014) and subsequent in silico normalization. Assemblies were annotated with the Trinotate annotation pipeline (Grabherr et al., 2011).

### Immune effector gene identification

Orthofinder 2.5.4 (Emms & Kelly, 2016) was used was to infer *T. molitor, C. vicina* and *P. apterus* orthologs of annotated immune genes from previously published insect genome projects (Benoit et al., 2016; dos Santos et al., 2015; Herndon et al., 2020; International Aphid Genomics Consortium, 2010; International Silkworm Genome Consortium, 2008). Additionally, blast and HMM homology searches were performed using previously described insect immune effector proteins as queries against each *de novo* assembly.

### Differential gene expression

Differential gene expression was determined using the R Bioconductor package DESeq2 v. 1.26.0 (Love, Huber, & Anders, 2014). For all three insect species, transcript abundances were quantified by pseudo-aligning RNAseq reads to *de novo* assemblies using Salmon v. 1.8.0 (Patro, Duggal, Love, Irizarry, & Kingsford, 2017). The R package tximport (Soneson, Love, & Robinson, 2015) was used to import salmons transcript-level quantifications into R v. 3.6.3 (R Core Team, 2020). The tximport object was used in conjunction with the sample metadata to make a DESeqDataSet object using the *DESeqDataSetFromTximport* function from the R package DESeq2 (Love et al., 2014). The DESeqDataSet object was used to identify differential expression as a function of developmental stage using the *DESeq* function. A likelihood-ratio test was used to compare a full model containing developmental stage as term to a reduced (intercept-only) negative binomial GLM to identify differentially expressed genes. All genes with a false discovery rate (FDR) corrected p-value less than 0.05 were considered differentially expressed genes. The mean of the normalised counts for each gene was used as the informative covariate for independent hypothesis weighting (Ignatiadis, Klaus, Zaugg, & Huber, 2016) to optimise the power of multiple testing.

The normalised counts were regularised log-transformed (rlog) for the PCA plots. The R package ggplot2 (Wickham, 2011) was used for plotting.

## Results

### Tenebrio molitor

A total of 13 differentially expressed immune effectors, including three lysozymes, nine AMPs and *prophenoloxidase*, a type-3 copper innate immunity protein, were identified in *Tenebrio molitor* during pupation. The AMPs included attacins (*attacin 1, attacin 2, tenecin 4),* one cecropin, coleoptericins (*coleoptericin A, coleoptericin B, tenecin 2*), the defensin *tenecin 1,* and the antifungal thaumatin *tenecin 3.* These immune effectors have been previously described by Johnston, Makarova, & Rolff (2014). Throughout *T. molitor* development, attacins and coleoptericins showed peak expressions at the onset of pupation (stage IV). The expressions of the immune effector at the six defined developmental stages in the last instar larva and pupa of *T. molitor* are shown in figure 1. The read count abundances, averaged over all developmental stages, of these differentially expressed immune effectors ranged from 60.53 (*I-type lysozyme 1*) to 19,962.58 (*Tenecin 1*) and had an overall mean of 8,509.60 (± 1,844.99 SE) read counts. A 14th identified immune effector, an attacin, was not differentially expressed (corrected p-value > 0.05) and had a mean read count of 158.44 (± 76.18 SE). The normalized read counts for the differentially and the non-differentially expressed immune effectors are shown in table 2 (supplement).

**Figure 1:**
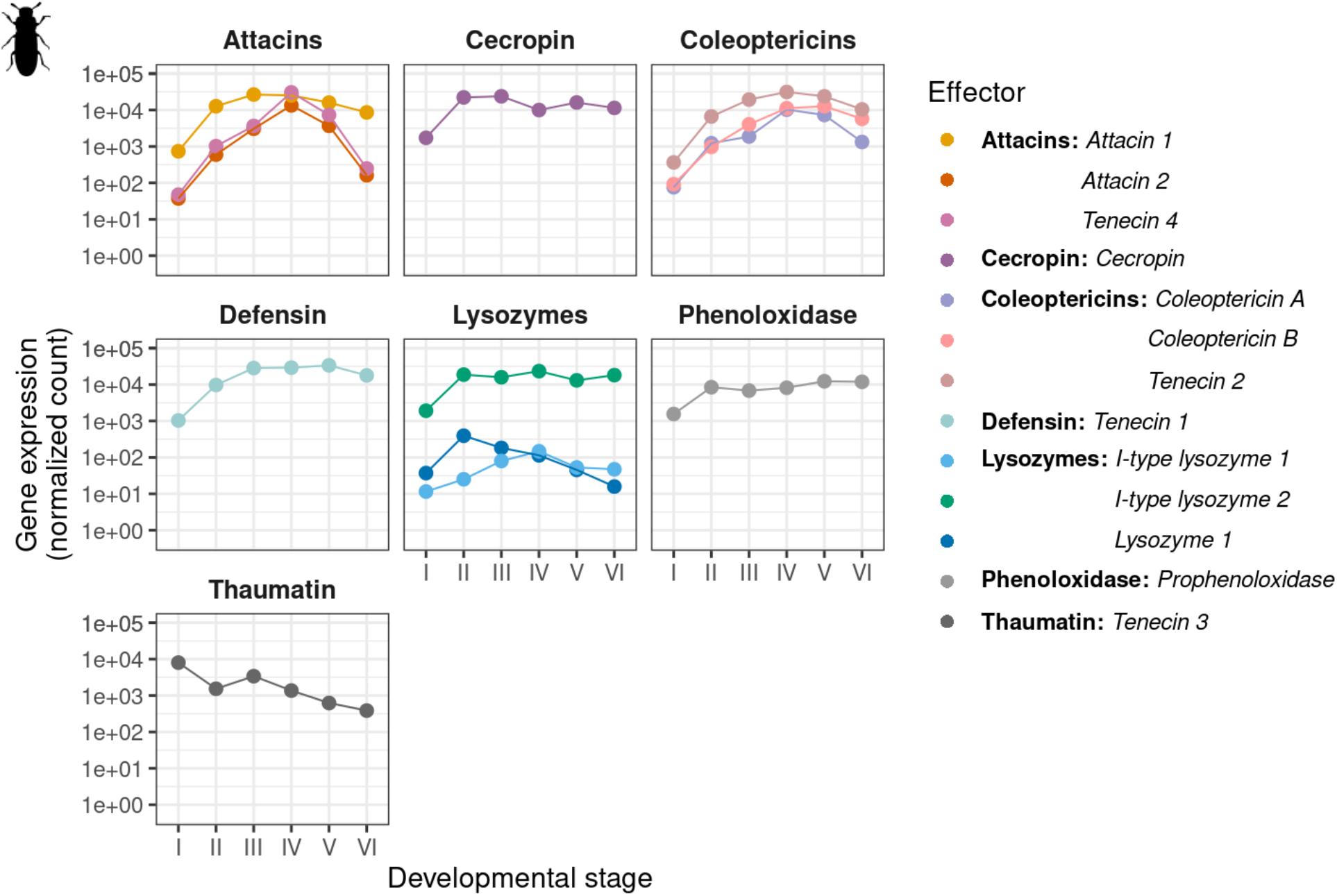
Differentially expressed immune effectors (corrected p-value < = 0.05) in the gut during the larval–pupal moult of *Tenebrio molitor.* Roman numerals correspond to specified developmental stages: (I) Last instar larvae, (II) little movement, (III) no movement, (IV) pupation after 1-6 h, (V) 12-20 h, and (VI) 24-48 h (see Material and Methods for more details). Plotted values represent the coefficients and 95% confidence intervals from negative binomial generalized linear models.

### Calliphora vicina

We identified six differentially expressed immune effector genes, which encoded two lysozymes and four AMPs, including two attacins and two diptericins. One lysozyme (lysozyme b) first increased in normalized read counts with peak read counts one hour after pupation in the fourth developmental stage and then decreased (see figure 2). The attacins also showed peak expressions one hour after pupation (stage IV). Figure 2 shows all six differentially expressed immune effectors of *C. vicina* at the six defined developmental stages in the last instar larva and pupa. The read count abundances, averaged over all developmental stages, of these differentially expressed immune effectors ranged from 17.45 (*diptericin a*) to 1,290.21 (*attacin a).* They had an overall mean of 668.21 (± 183.58 SE) read counts. Lysozymes, attacins and diptericins have been previously described in *C. vicina.* Diptericin has been shown to be released by the blowflies’ hemocytes (Gordya et al., 2017; Yakovlev et al., 2017). Yoon et al. (2022) showed high expression levels of attacin c, diptericins and lysozymes in the final instar larvae of *C.vicina.*

**Figure 2:**
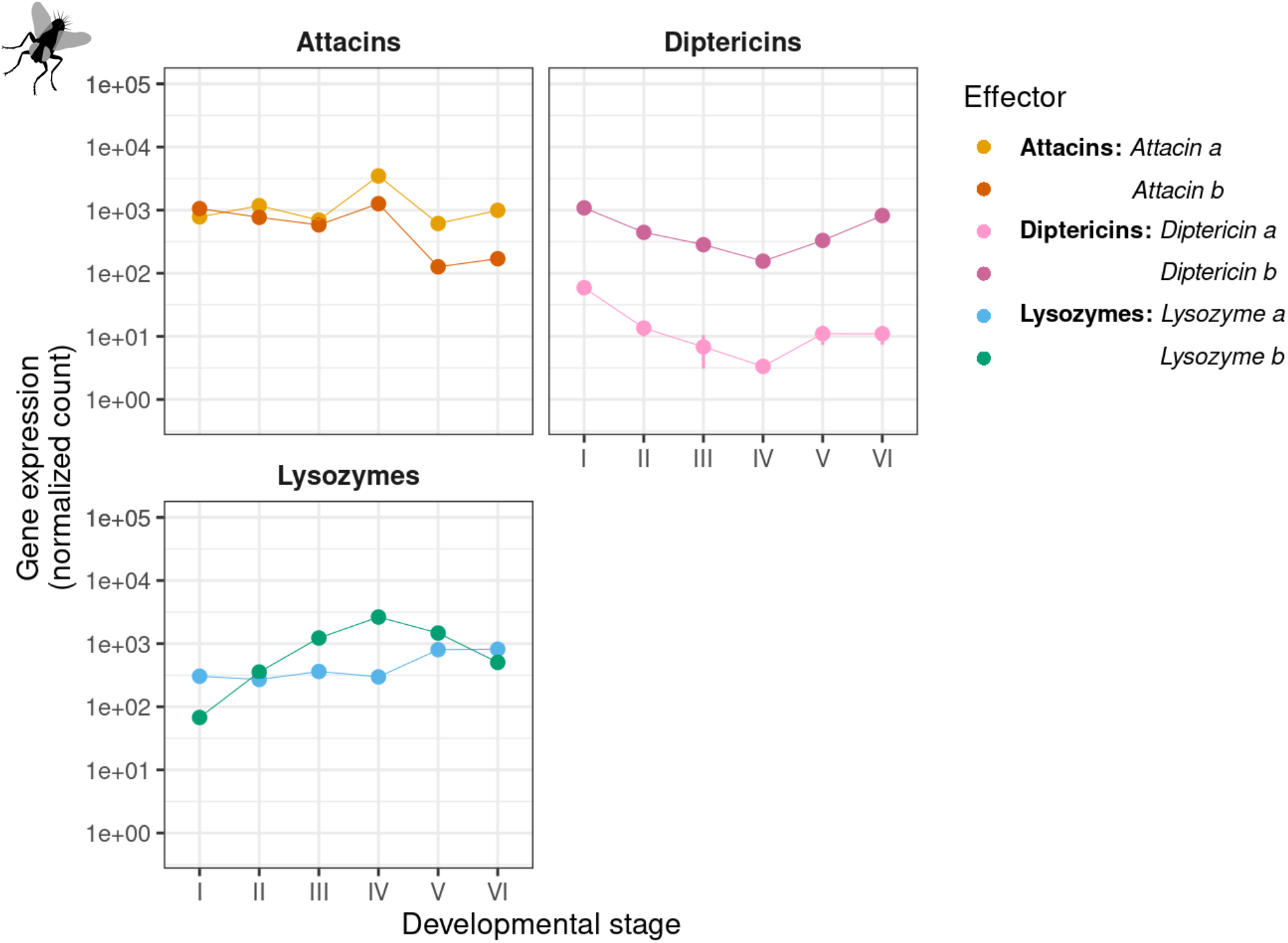
Differentially expressed immune effectors (corrected p-value <= 0.05) in the gut during the larval–pupal moult of *Calliphora vicina.* Roman numerals correspond to specified developmental stages: (I) Post feeding larvae, (II) pre-pupal stage, (III) pupal stage at the onset, (IV) pupae after 1 h, (V) 4-6 h, and (VI) 8-12 h (see Material and Methods for more details). Plotted values represent the coefficients and 95% confidence intervals from negative binomial generalized linear models.

A total of 30 identified immune genes had a corrected p-value greater than 0.05 and were not differentially expressed. These genes encoded six types of immune effectors: lysozymes, attacins, coleoptericins, diptericins, cecropins and defensins. The read count abundances, averaged over all six developmental stages, of these non-differentially expressed immune effectors ranged from 0.27 (attacin j) to 691.35 (attacin a) and had an overall mean of 72.22 (± 28.71 SE) read counts. Table 3 in the supplement gives an overview of the normalized count means for the differentially and non-differentially expressed immune effectors and the number of unique immune effectors within each effector group.

### Pyrrhocoris apterus

A total of nine immune effectors were identified in the *P. apterus de novo* assembly, of which three (*lysozyme*, *c-type lysozyme* and *phenoloxidase*) were differentially expressed at very low normalised read count abundances ranging from 8.15 (*Lysozyme*) to 41.21 (*C-type lysozyme*) and with an overall mean of 26.20 (± 9.66 SE) read counts (averaged over all five developmental stages). The lysozymes showed highest expressions one day before the larval-adult moult (stage III) and phenoloxidase four to five days after the moult into the final instar (stage II). Figure 3 shows the expressions of these immune effectors throughout the five defined developmental stages in the last instar larva and adult. The other immune effectors consisted of three I-type and two C-type lysozymes. Supplementary table 4 gives an overview of the normalised read count abundances for the differentially and non-differentially expressed immune effectors.

**Figure 3:**
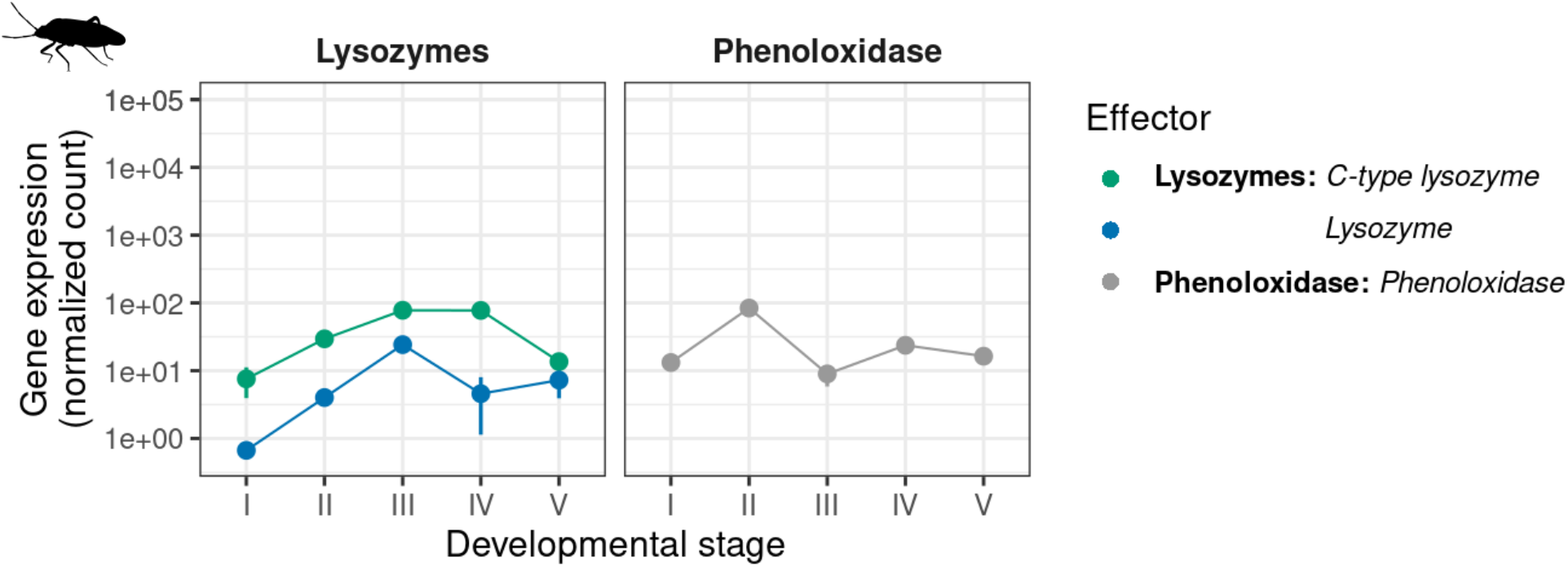
Differentially expressed immune effectors (corrected p-value <= 0.05) in the gut during the larval–adult moult of *Pyrrhocoris apterus.* Roman numerals correspond to specified developmental stages: (I) Freshly moulted last instar larvae, (II) last instar larvae after 4-5 d, (III) 9 d and (IV) 9.5 d, and (V) freshly eclosed adults (see Material and Methods for more details). Plotted values represent the coefficients and 95% confidence intervals from negative binomial generalized linear models.

## Discussion

We found high read count abundances of differentially expressed immune effectors in the gut at the larval-pupal moult of the two holometabolous insects, *Tenebrio molitor* and *Calliphora vicina;* no such high abundances were observed at the larval-adult moult in the hemimetabolous insect *Pyrrhocoris apterus*. We also found peak expressions of immune effectors at the onset of pupation of the two holometabolous insects*. P.apterus* showed the highest expressions of immune effectors before the larval-adult moult. Our findings of high immune effector expressions in holometabolous but not hemimetabolous insects during metamorphosis are consistent with Johnston et al. (2019) and confirm that only complete metamorphosis seems to elicits a prophylactic gut immune response as an adaptive response in holometabolous insects, which controls the microbiota during gut replacement (Russell & Dunn, 1991, 1996).

The immune effectors identified in the mealworm beetle *T. molitor* have been previously described by Johnston et al. (2014) as components of the mealworm beetle’s immune system. They identified immediate and long-lasting immune responses in the course of seven days after an immune challenge with heat-killed bacteria. The immediate response included the upregulation of phenoloxidase, which produces cytotoxic melanin and oxidative intermediates with broadspectrum antibacterial activity (Zhao et al., 2007). In our study, phenoloxidase was differentially expressed and increased in the level of expression from the first (freshly moulted final instar larvae) to the second developmental stage (little movement stage, 1-2 d before pupation). It then did not change in the expression level throughout the remaining developmental stages of our study. The long-lasting induced immune effectors identified by Johnston et al. (2014) were antibacterial peptides and the iron-sequestering protein ferritin. These AMPs included attacins, coleoptericins and the defensin *tenecin 1.* Other AMPs, including cecropins, the attacin *tenecin 4,* the coleoptericin *tenecin 2* and the antifungal thaumatin *tenecin 3,* as well as lysozymes, were not upregulated after the immune challenge. We found peak expressions of *tenecin 2* and *tenecin 4* at the onset of pupation, indicating a prophylactic gut immune response.

The blowfly *C. vicina* showed peak expressions of antimicrobial proteins and peptides in the brown pupa (stage IV). Nunes et al. (2021) found in another dipteran species, the fruit fly *Drosophila melanogaster,* peak expressions of three AMPs (*drosomycin, drosomycin-like 2* and *drosomycin-like 5*) in the motionless white pre-pupa and discussed these AMP peaks as a prophylactic immune response. The AMP *drosomycin-like 2* was recruited in the midgut (Nunes et al., 2021), which undergoes extensive remodelling at complete metamorphosis (Martín-Vega, Simonsen, & Hall, 2017).

In the firebug *P. apterus*, we found lysozymes and phenoloxidase upregulated with very low read count abundances compared to the peak expressions of the two holometabolous insect species. The less drastic changes of the gut of hemimetabolous insects during metamorphosis, which does not entail gut replacement (Teixeira, Fialho, Zanuncio, Ramalho, & Serrão, 2013), are likely to explain the differences. The more minor changes during incomplete metamorphosis seem not to necessitate prophylaxis. The lysozymes in *P. apterus* were strongest upregulated one day (stage III) and 12 hours before adult eclosion (stage IV) and phenoloxidase four to five days after the moult in the final instar (stage II). Phenoloxidase is a central enzyme secreted by the salivary glands of phytophagous Hemiptera, including Pyrrhocoridae (Hori, 2000). Insect saliva has numerous functions, including digestion and antimicrobial activity. According to Miles (1969), polyphenol oxidase seems to be an invariable component of the watery saliva of phytophagous Hemiptera. It serves as a counter to defensive toxins in the insects’ food (Miles, 1969). Also, the immune system of insects can control beneficial symbionts (Login et al., 2011). In another species from the Pyrrhocoridae family, the cotton stainer bug *Dysdercus fasciatus,* beneficial and heritable gut bacterial symbionts induce an immune response with an upregulation of *c-type lysozyme* and the AMP*pyrrhocoricin* (Bauer, Salem, Marz, Vogel, & Kaltenpoth, 2014).

However, experimental support for a prophylactic effect in holometabolous insects is lacking. We also cannot exclude the possibility that in either species, there may be true differentially expressed immune genes that were not successfully annotated. Johnston et al. (2019) discuss that alternatively to a prophylactic effect, immune induction may serve to control the proliferation of the microbiota as they observed that the upregulation of immune effectors persisted into the pupa when the gut of their studied greater wax moth *Galleria mellonella* undergoes apoptosis and necrosis to release breakdown products that are recycled by the replacement gut. This indicates the possibility that immune induction may suppress bacterial growth that otherwise disrupts the complex trophic relationship between the autolytic larval and the replacement adult gut. A second alternative explanation, Johnston et al. (2019) discuss, is that the observed immune induction drives changes in microbial community composition, facilitating ontogenetic habitat and/or diet shifts.

## Acknowledgements

We are grateful to Elisa Bittermann, Luisa Linke, Jia Clara Kim and Marlene Finger for support in the lab. This research was funded by the DFG (Deutsche Forschungsgemeinschaft, Grant RO 2284/4-1).

## Appendix

**Table 1:**
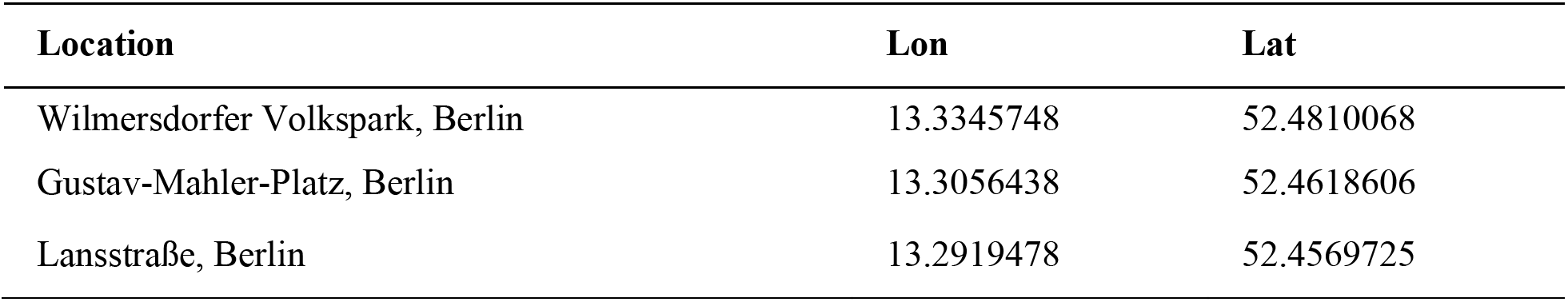
Sampling locations (coordinates) of the *Pyrrhocoris apterus* from *Tilia cordata* trees in Berlin.

**Table 2:**
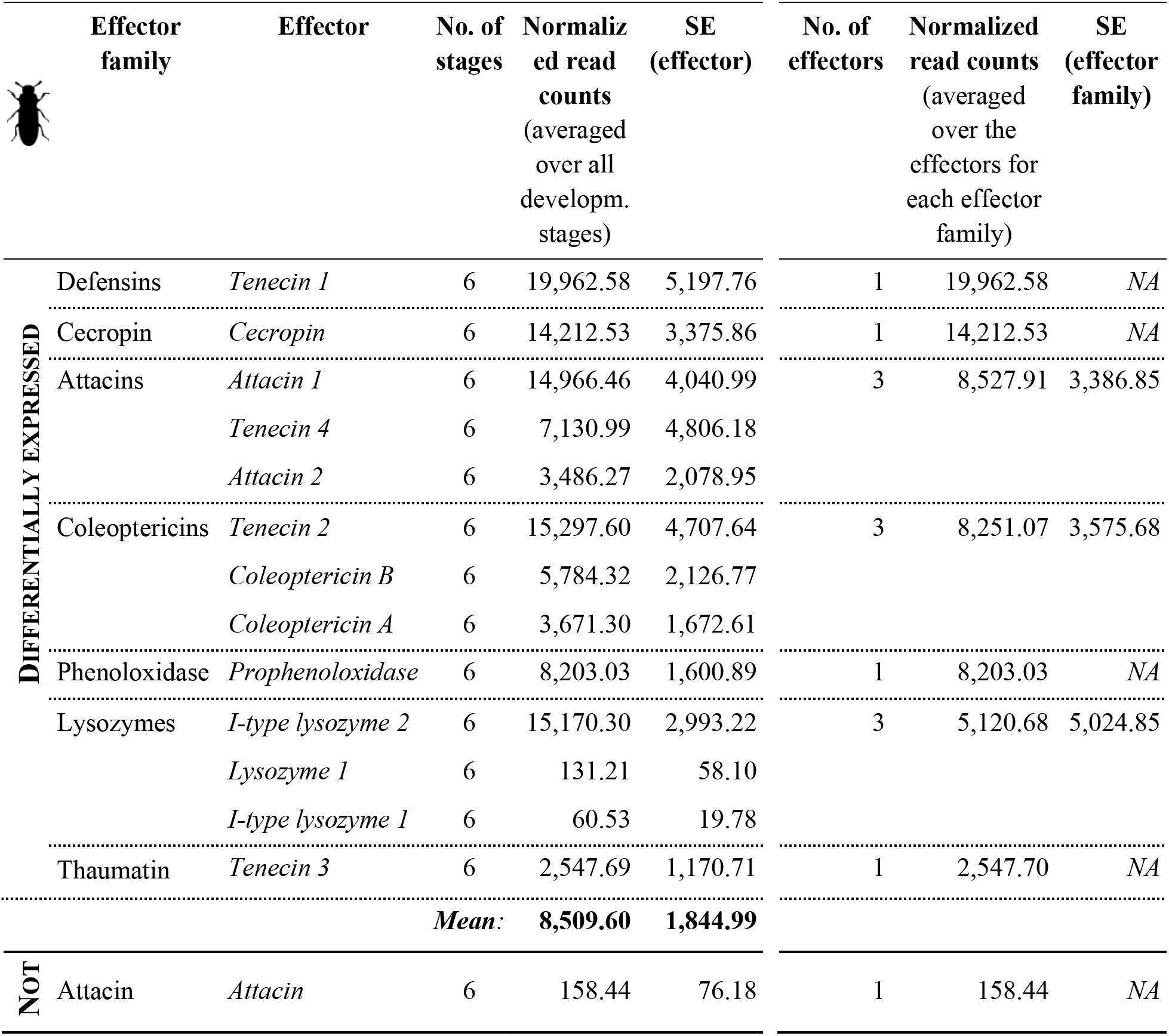
Shown are the immune effectors of *Tenebrio molitor* and their normalized read counts for each differentially (corrected p-value <= 0.05) and non-differenially expressed (corrected p-value > 0.05) immune effector (averaged over all six developmental stages). Also shown are the read count means for each effector family and the overall mean.

**Table 3:**
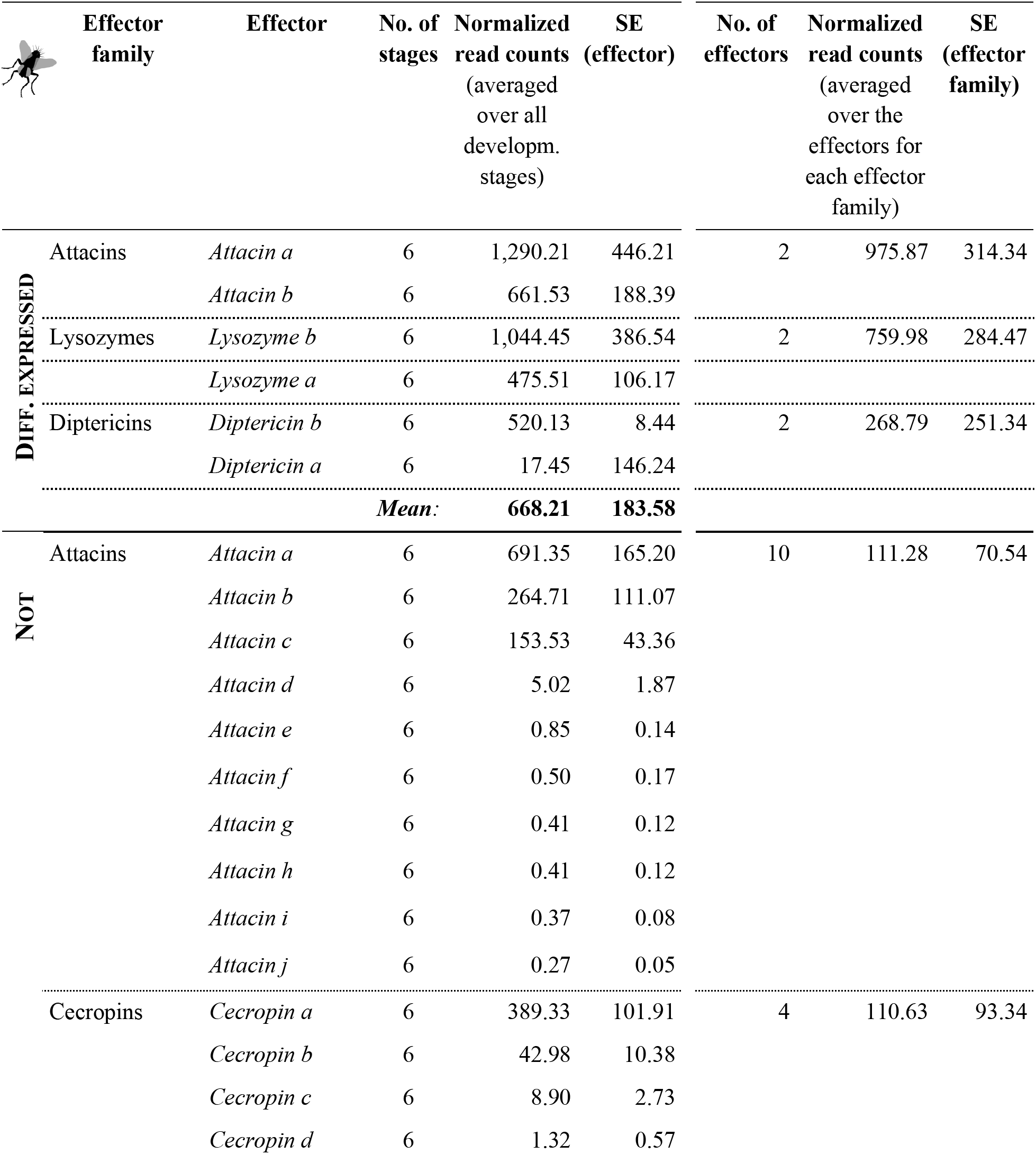
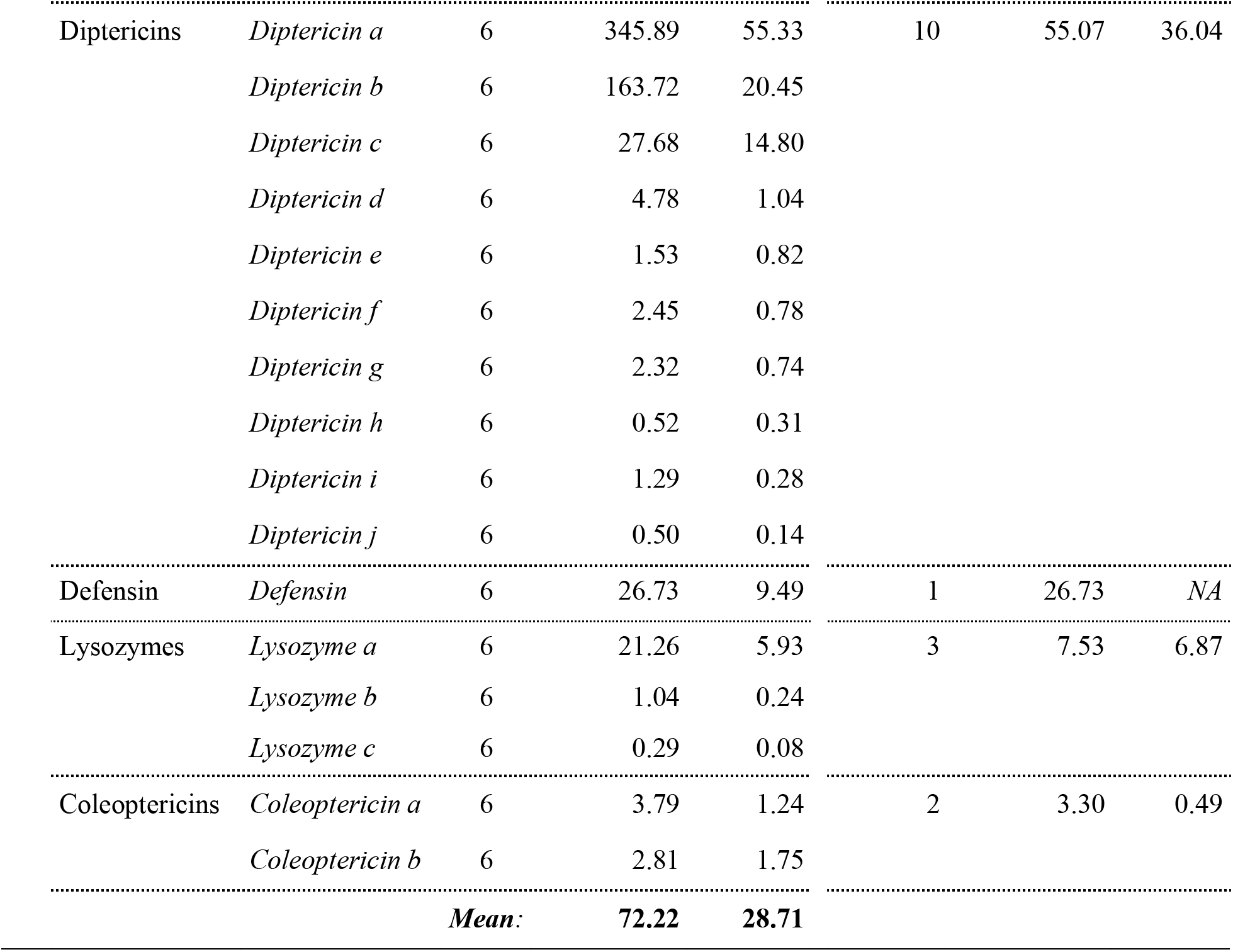
Shown are the immune effectors of *Calliphora vicina* and their normalized read counts for each differentially (corrected p-value <= 0.05) and non-differenially expressed (corrected p-value > 0.05) immune effector (averaged over all six developmental stages). Also shown are the read count means for each effector family and the overall mean.

**Table 4:**
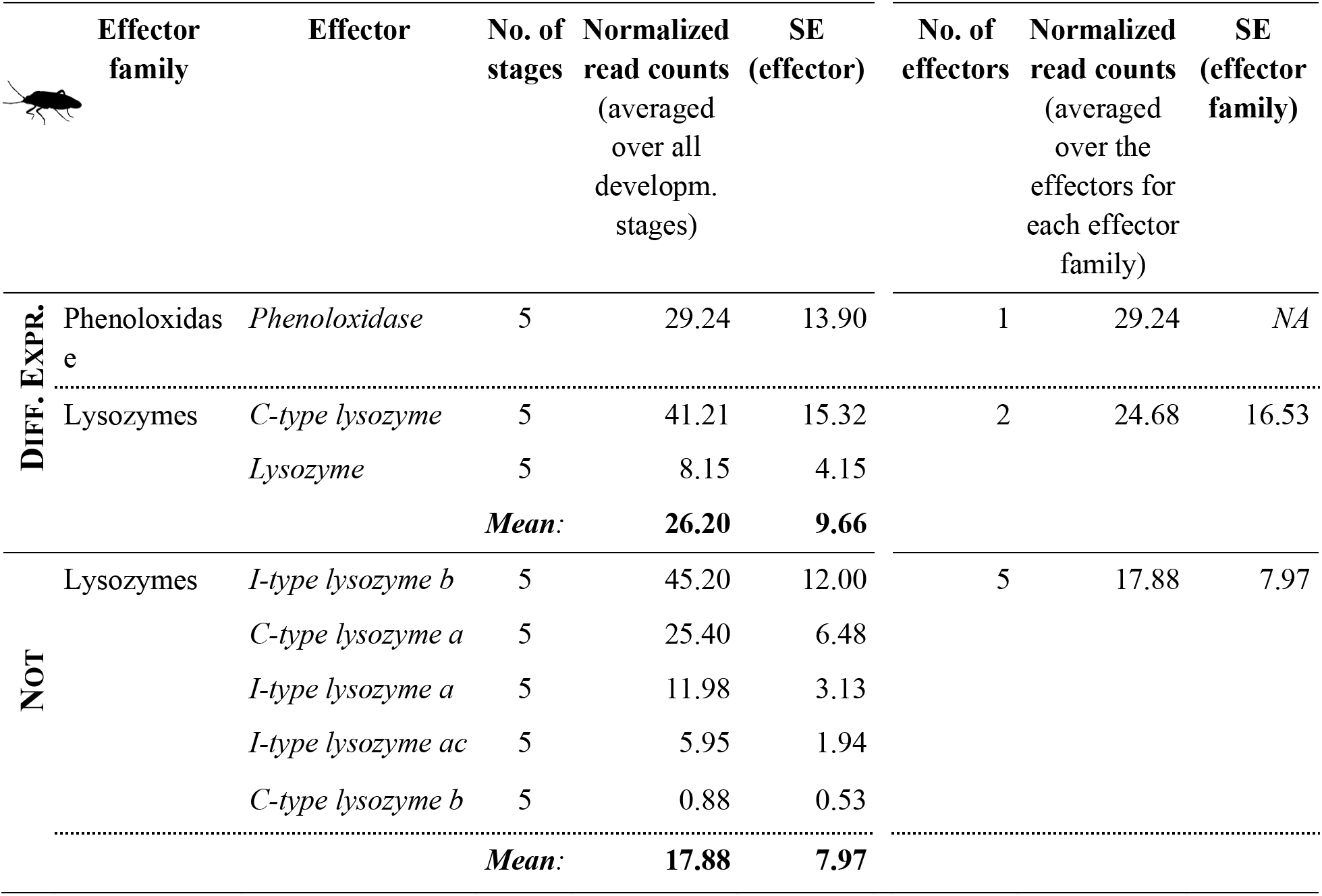
Shown are the immune effectors of *Pyrrhocoris apterus* and their normalized read counts for each differentially (corrected p-value <= 0.05) and non-differenially expressed (corrected p-value > 0.05) immune effector (averaged over all six developmental stages). Also shown are the read count means for each effector family and the overall mean.

**Figure S4:**
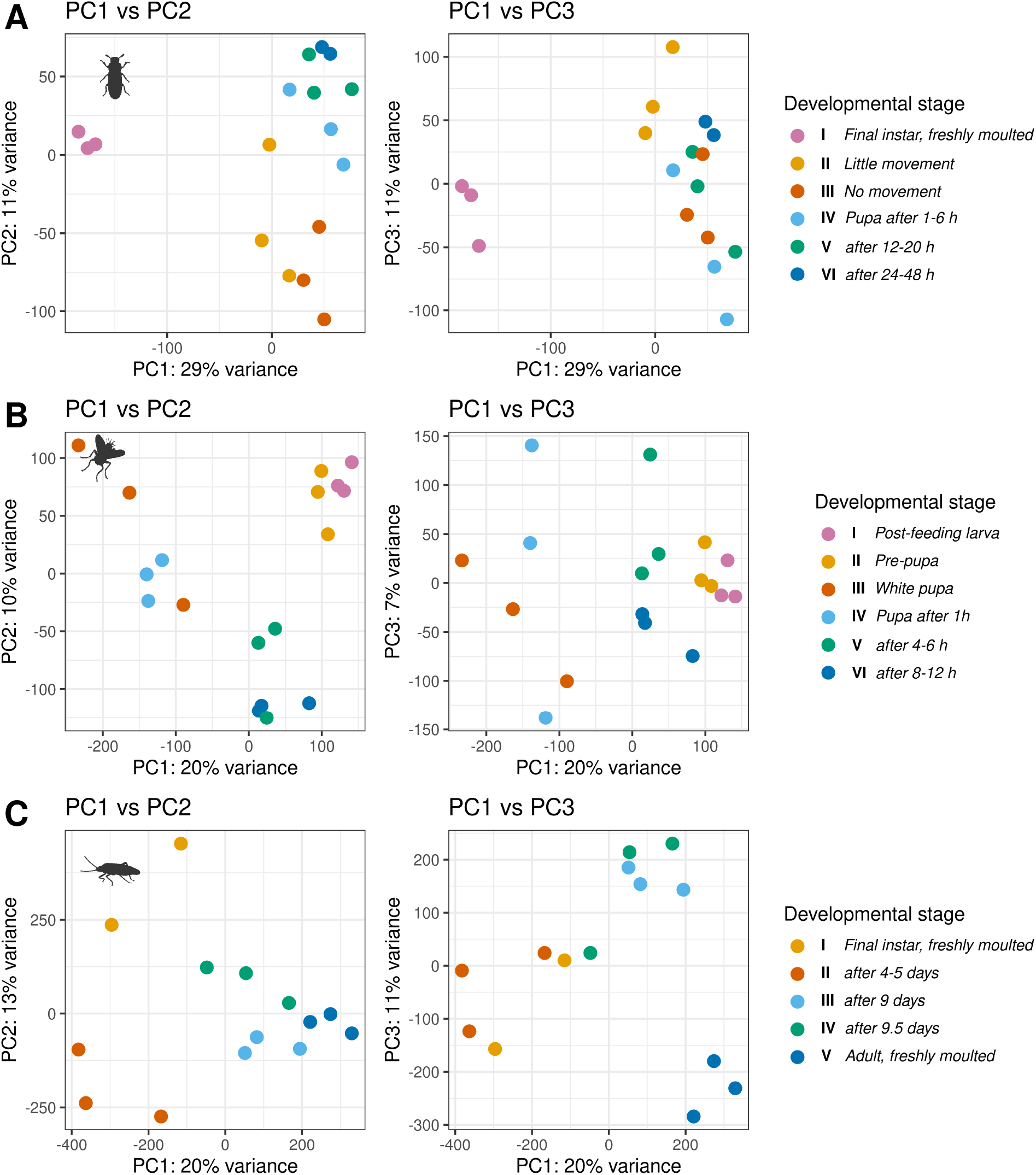
Principle components analysis (PCA) of *Tenebrio molitor* (A), *Calliphora vicina* (B) and *Pyrrhocoris apterus* (C) for examining the relationships between the developmental stages.

## Notes

### Competing Interest Statement

The authors have declared no competing interest.

### Summary of Updates

a few typos

